# Genetic Bases Of Aposematic Traits: Insights from the Skin Transcriptional Profiles of *Oophaga* Poison Frogs

**DOI:** 10.1101/706655

**Authors:** Andrés Posso-Terranova, José Andres

**Author notes:** All authors contributed equally to this work.

## Abstract

Aposematic organisms advertise their defensive toxins to predators using a variety of warning signals, including bright coloration. While most Neotropical poison frogs (Dendrobatidae) rely on crypsis to avoid predators, *Oophaga* poison frogs from South America advertise their chemical defenses, a complex mix of diet-derived alkaloids, by using conspicuous hues. The present study aimed to characterize the skin transcriptomic profiles of the South American clade of *Oophaga* poison frogs (*O. anchicayensis, O. solanensis, O. lehmanni* and *O. sylvatica*). Our analyses showed very similar transcriptomic profiles for these closely related species in terms of functional annotation and relative abundance of gene ontology terms expressed. Analyses of expression profiles of *Oophaga* and available skin transcriptomes of cryptic anurans allowed us to propose possible mechanisms for the active sequestration of alkaloid-based chemical defenses and to highlight some genes that may be potentially involved in resistance mechanisms to avoid self-intoxication and skin coloration. In doing so, we provide an important molecular resource for the study of warning signals that will facilitate the assembly and annotation of future poison frog genomes.

## Introduction

Aposematic organisms advertise their defensive toxins to predators using a variety of warning signals [1–3]. Among aposematic species, the poison frogs (Dendrobatidae) from the tropical rain forests of Central and South America represent one of the most spectacular examples of warning coloration. While the majority of dendrobatids rely on crypsis to avoid predators, some members of this family are both brightly colored and chemically defended [4]. From a genetic point of view, these aposematic defenses can be defined as a complex phenotype resulting from the integration (*i*.*e*. covariation) of different genetic elements related to conspicuousness, bold behavior, unpalatability, diet specialization, etc. While a long history of research has been devoted to understand the genetics of warning coloration in arthropods, particularly in *Heliconius* butterflies [5, 6], the molecular underpinnings of aposematism in vertebrates, particularly the mechanisms whereby individuals become toxic (or distasteful) remain mostly unknown. In this study, our primary aim was to characterize the skin transcriptomic profiles of *Oophaga* species as a first attempt to shed light on the molecular genetic bases of aposematic components in poison frogs: the ability to sequester alkaloid-based chemical defenses, warning coloration, and the resistance mechanisms to avoid self-intoxication.

The frogs of this complex inhabit the lowland Pacific rainforests of the Colombian and Ecuadorian Chocó. Preliminary molecular data from nuclear and mitochondrial markers showed a similar genetic background among lineages in contrast to an extraordinary diversity of morphotypes [7]. Individuals from different allopatric lineages can be relatively homogeneous, striped, or spotted, and their colors range from bright red, to orange and yellow (Fig 1) [8]. These polymorphic coloration patterns serve as a warning signal of their chemical defenses [9], a complex mix of diet-derived alkaloids secreted by the dermal glands [10, 11]. Although these chemicals may involve metabolism and transport through other tissues, the final accumulation of toxins as well as the production of pigmentary cells is performed in the skin tissue of these organisms even during the adult stage [12–14]. Thus, we expected that the genes, pathways, and/ or gene networks potentially associated with coloration, alkaloid metabolism, transport and storage, to be expressed in the skin of these lineages.

**Fig 1.**
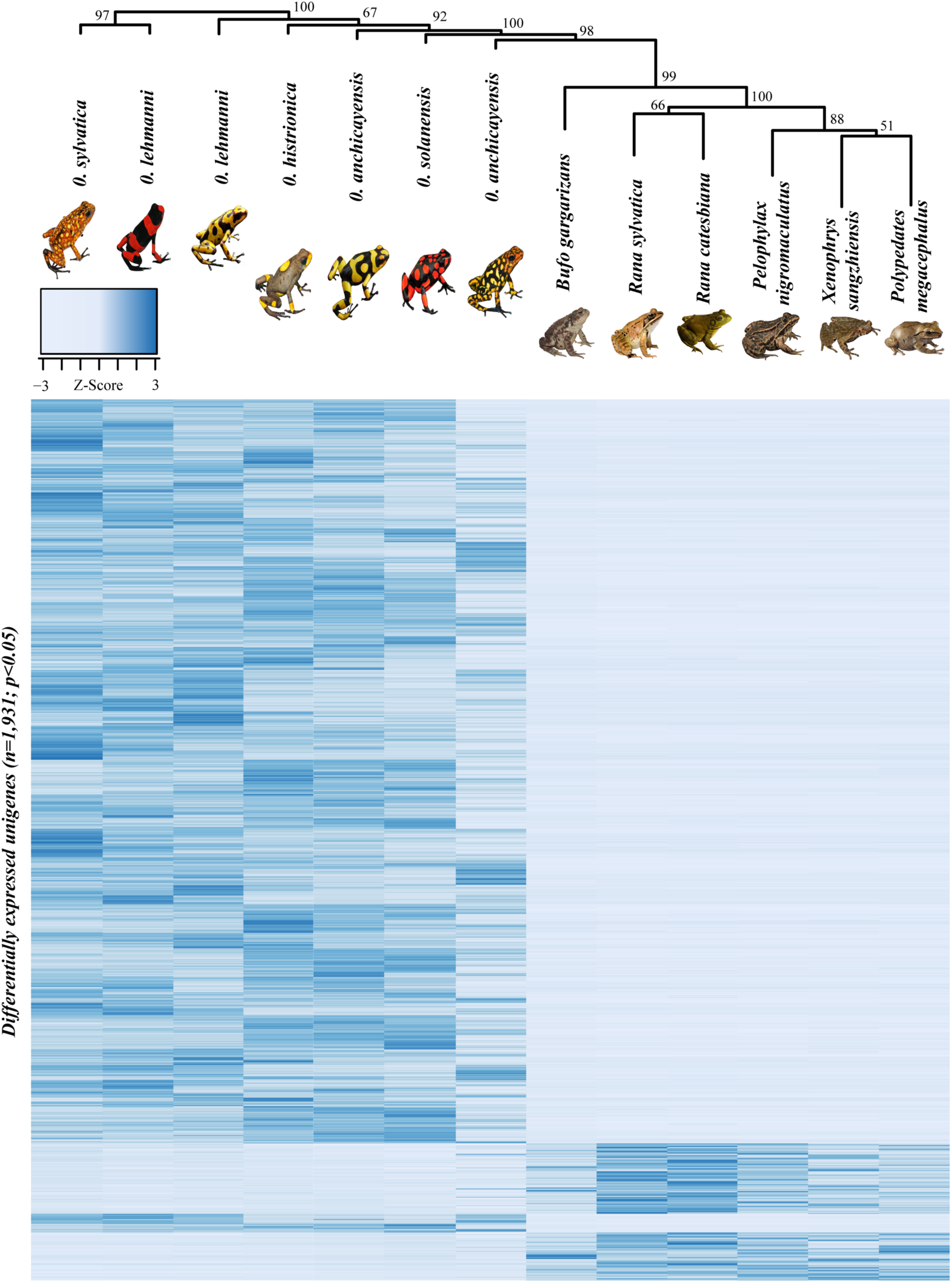
Hierarchical clustering (bootstrap=999) and heatmap contrasting the differentially expressed unigenes from skin tissue between *Oophaga* and cryptic species. Color within the heatmap represent the log2 fold change values. Dark blue indicates upregulation while a lighter coloration indicates downregulation. Rows of the heatmap correspond to the differentially expressed unigenes (n=1,931) detected in this study.

Although the lineages of the *Oophaga* studied here display a wide variety of warning signals, most of them share a black background coloration (Fig 1). In adult frogs, this melanistic coloration is controlled by the lack of iridophores and xanthophores in the dermal chromatophore unit, as well as the size and dispersion pattern of the melanin-containing organelles (melanosomes) within the dermal melanophores [15]. Thus, we also hypothesized that in adult individuals, genes involved in the amount, size and distribution of the melanosomes should be differentially expressed in the skin. Similarly, because the secreted alkaloids of chemically-defended dendrobatids are known to disrupt the normal ion-channel activity and to alter the neurotransmitter-receptor binding capacity of nerve and muscle cells [16–18], we hypothesized that the fraction of genes differentially expressed in the skin of poison frogs (when compared to non-aposematic species) is also enriched in genes related to alkaloid sequestration, transport and autotoxicity avoidance.

To test these hypotheses, we first assembled the skin transcriptomes of five different Harlequin poison frogs species: *O. histrionica* [19], *O. lehmanni* [20], *O. sylvatica*, [21] and the recently described *O. anchicayensis* and *O. solanensis* [22]. Then, we compared the transcription profiles of these aposematic species with that of a diverse group of cryptic anurans. Functional annotation of unigenes showing a phylogenetic signal (*i*.*e*. highly represented and differentially expressed in *Oophaga* poison frogs) allowed us to propose plausible metabolic pathways (and candidate genes) related with toxicity and color. Overall, this study provides an important molecular resource for the study of aposematism within poison frogs and will facilitate the future assembling and annotation of the complex dendrobatid genomes [23].

## Methods

### Library preparation and sequencing

Due the conservation status of these frogs (CR, EN, VU; IUCN) and the restrictions imposed by the Ethics Committee, we were allowed to euthanize a very limited number of samples per species. Seven individuals (*O. lehmanni* and *O. anchicayensis*, n=2 per each species; *O. histrionica, O. sylvatica*, and *O. solanensis*, n=1; each species) were euthanized in the field with benzocaine gel at 5% by direct application into the mouth cavity. Individual skin samples were taken and stored at −80 °C in RNAlater^®^ (Life Technologies, Carlbad, CA). RNA extractions were performed using Tri-Reagent (Molecular Research Center, Cincinnati, OH). After quality check (Agilent Bioanalyzer Agilent Technologies, Wilmington, DE) samples were used to generate RNA-seq libraries using the Illumina Truseq RNA Sample Prep protocol (Illumina, San Diego, CA). Libraries were cleaned using AMPure XP and sequenced on a single Illumina HiSeq2000 lane (TruSeq SBS v. 3) as follows: four libraries (*O. anchicayensis, O. histrionica, O. lehmanni* and *O. sylvatica*) were sequenced using single-end (150-bp) while the remaining three libraries (*O. anchicayensis, O. solanensis* and *O. lehmanni*) were run in a single lane of a paired-end module (100-bp, × 2). All animal procedures were approved by the Ethics Committee of Universidad Nacional de Colombia (Acta No.03, July 22^nd^, 2015) and were conducted based on the NIH Guide for the Principles of Animal Care. Sampling was conducted according to the research permits granted by Autoridad Nacional de Licencias Ambientales ANLA, Resolución 0255 del 12 de Marzo de 2014 and Resolución 1482 del 20 de Noviembre de 2015. Sequencing was carried out by the Cornell University’s BioResource Center. Datasets including raw RNA sequence reads are available from the authors upon request.

### Transcriptome assemblies and functional annotation

Initial read quality trimming, filtering and removal of adapters was performed using *FLEXBAR* v.2.5 [24]. All retained reads were ≥ 50bp with an average quality of ≥ 30 and less than two uncalled (ambiguous) bases (S1 Table). For each library, a *de novo* transcriptome was assembled using *TRINITY* v.2.1.0 [25] with default settings (kmer=25, minimum contig length=48) and keeping only the longest transcript per cluster for subsequent analyses. Varying these parameters (i.e. default settings) did not result in assemblies with longer N50 values. To account for differences in the quality and sequencing strategy among individual transcriptomes, we constructed a composite *de novo* reference transcriptome. Briefly, we followed [26] and combined 20% randomly selected reads (>33 million in total) from each of the four single-end libraries (*O. anchicayensis, O. histrionica, O. lehmanni* and *O. sylvatica*). Then, a *TRINITY* assembly was performed using a minimum contig length of 350 bp. To filter out highly similar contigs that may potentially represent alternatively spliced transcripts, we implemented the error correction module of *iAssembler* v1.3.2 [27] with default parameters (maximum length of end clips=30 bp, minimum overlap length=40 bp, minimum percent sequence identity=95%).

Gene annotation was conducted using a sequential *BLASTX* search to both, the available *Xenopus* transcriptomes and the NCBI non-redundant (*nr*) database. The composite transcripts were first compared with that of the *Xenopus* databases retaining annotations with E-values ≤ 10^−5.^ Unannotated contigs were then submitted for *BLASTX* to the *nr* protein database for possible identification. Gene ontology (GO) annotation and term mapping was performed using *Blast2Go* v 5.2.0 with default significance cutoffs [28].

To estimate the completeness of each transcriptome, we implemented the expected gene content of Benchmarking Universal Single-Copy Orthologs (*BUSCO* v3) [29] as implemented in *gVolante* v1.1.0 [30]. This widely used metrics for transcriptome assembly [31] assesses the quality of any given transcriptome by estimating the completeness of core vertebrate genes predicted to be ubiquitous in eukaryotes. The pre-processing of the reads and all *BLASTX* analyses were run in the Bugaboo Dell Xeon cluster of the western Canada’s WestGrid computer facilities (www.westgrid.ca).

### Transcriptome profiles

Analyses on the distribution of reads and the relative abundance of different transcripts within each individual sample (*i*.*e*. lineage) were carried out by mapping RNA-seq reads back to the composite *de novo* assembly. In order to identify highly represented unigenes, we first implemented eXpress (probabilistic assignment of ambiguously mapping sequenced fragments) [32] to estimate the effective number of reads (ER) that mapped to the contigs in the reference transcriptome after adjusting for read number and length biases. Then, to compare the proportion of reads that mapped to a transcript in each *Oophaga* RNA-library, we estimated the number of transcripts per million (TPM), a measure of RNA abundance that allows the comparison between samples (sum of all TPMs in each sample are the same) [33]. We visually explored for highly represented unigenes in each RNA-library by using dispersion plots of the ER values and percentile plots of the TPM distribution. Additionally, we performed a Principal Component Analysis (PCA) of the reference contigs dataset and using each library TPMs values as independent variables and selected the overall top expressed unigenes (2%).

To further characterize the skin transcriptome profile of *Oophaga* poison frogs, we compared the transcription levels of annotated unigenes in this clade with those observed in several cryptic (*i*.*e*. non-aposematic) anurans, including a toad (*Bufo gargarizans*), and five frog species *(Pelophylax nigromaculatus, Polypedates megacephalus, Rana catesbeiana, Rana sylvatica*, and *Xenophrys sangzhiensis*, S2 Table) [34, 35]. RNA-seq data from these species was obtained from the Sequence Read Archive platform (SRA) and represented the only skin-based RNA-seq available to us for comparison purposes. Although transcriptomic datasets are available for phylogenetically closer and cryptic species (i.e *Colostethus*) [36], these datasets were not included in our analysis because they are not derived from skin tissue and may bias our analyses. Read quality, contamination screening, duplicate removal, and quality trimming steps were performed in FLEXBAR v.2.5 as previously described. The filtered raw reads from individual libraries (*Oophaga*, n=7; cryptic species n=6) were mapped to our composite *de novo* reference transcriptome using *Bowtie2* v2.3.4.1 [37]. To accommodate for sequence divergence among the taxa included in our study (≈75–98 similarity), we followed [38] and ran the alignment algorithm allowing for soft clipping (*--local*) and with a maximum penalty value of three (*--mp 3*). Then, we implemented *eXpress* [32] to estimate the effective number of reads (ER) that mapped to the contigs in the reference transcriptome after adjusting for read number and length biases. Only unigenes with an average of >10 ER/library were included in the differential expression analyses [39].

To identify genes showing a phylogenetic signal in their expression levels and to estimate fold changes in expression levels between aposematic (*i*.*e. Oophaga*) and cryptic species, we followed [40] and implemented the R package *DEseq* [41] to select unigenes with an adjusted p-value of <0.05. To visualize patterns of expression, we constructed a multidimensional scaling plot (Euclidean distances) of the ER data using *Past* v. 3.18 [42]. A heatmap of the expression differences between *Oophaga* and cryptic anuran lineages (n_genes_=1,931) was generated using the ‘*heatmap*.*2*’ function of the ‘*gplots’* package, with Euclidean distances and complete linkage for clustering (bootstrap=999) [43]. *BLAST* analysis and GO term mapping for the 1,931 unigenes included in this analysis was performed as described above.

## Results

### Transcriptome assemblies and functional annotation

Illumina sequencing produced an average number of reads per sample of 118.3 million for paired-end and 37.7 million for single-end libraries. After a stringent read trimming involving the removal of low-quality sequences, duplicated reads and reads containing adapter/primer sequences, we retained over 81% of the initial sequencing data. Paired-end libraries produced a fairly consistent number of reads (S1 Table).

After the removal/merging of highly similar contigs (paired-end:7,5% - 9,6%; single-end: 6.0% - 13.4%), the 150-bp based transcriptomes recovered a large number of contigs ranging from 35,287 (*O. anchicayensis*) to 107,381 (*O. sylvatica*). Paired-reads libraries (100-bp; 2X) generated transcriptomes with a relatively similar number of contigs (40,398 for *O. solanensis* and 60,494 for *O. lehmanni*). N50 values and average transcript length (AL) were lower for paired-end libraries (N50= 498-561bp; AL= 441.78±379 - 475.93±436bp) than for assemblies produced with single-end libraries (N50=667-1579bp; AL=538.9±1001bp - 668.6±819 bp) (Table 1). Assembled and annotated transcriptomes are available from the authors upon request.

**Table 1.** Summary statistics and quality assessment estimators of the eight *de novo* assembled transcriptomes generated in this study (*Oophaga* species n=7; composite reference n=1).

|  | Paired-end libraries |  |  | Single-end libraries |  |  |  |  |
| --- | --- | --- | --- | --- | --- | --- | --- | --- |
|  | *O. solanensis* | *O. anchicayensis* | *O. lehmanni* | *O. sylvatica* | *O. lehmanni* | *O. anchicayensis* | *O. histrionica* | Reference |
| File size (Mbytes) | 22.23 | 32.06 | 35.7 | 93.9 | 77.3 | 23.16 | 85.97 | 31.6 |
| # contigs | 43,690 | 63,520 | 66,937 | 123,916 | 117,082 | 37,552 | 111,823 | 31,498 |
| # contigs after filtration<br>(removal/merging of<br>highly similar contigs) | 40,398 | 57,516 | 60,494 | 107,381 | 104,523 | 35,287 | 96,892 | 31,498 |
| # bases | 19,853,219 | 28,061,593 | 31,856,993 | 81,266,530 | 66,367,430 | 20,238,091 | 74,768,159 | 31,968,628 |
| # bases after filtration | 17,164,733 | 23,756,585 | 26,287,312 | 58,224,462 | 50,845,151 | 15,278,290 | 53,582,406 | 31,968,628 |
| Merged contigs | 7.5% | 9.5% | 9.6% | 13.3% | 10.7% | 6.0% | 13.4% | 0.0% |
| Average transcript<br>length (AL) | 454.41± 405 | 441.78± 379 | 475.93 ± 436 | 655.8 ± 799 | 566.8± 648 | 538.9 ± 1579 | 668.6 ± 819 | 1,014.91 ± 948.43 |
| Maximum length | 6,115 | 7,915 | 8,400 | 15,102 | 12,906 | 54,265 | 15,136 | 16,041 |
| N50 | 528 | 498 | 561 | 1079 | 809 | 667 | 1,101 | 1,316 |
| Contigs with *BLAST* hits | 46.5% | 47.7% | 50.8% | 42.1% | 41.6% | 53.4% | 44.5% | 62.6% |

Comparative *BLAST* analysis indicated that our transcriptomes recovered a significant proportion of the *X. laevis* and *X. tropicalis* transcriptomes (50,592 and 41,042 unigenes respectively). Paired-end transcriptomes showed a lower number of significant *BLAST* hits (18,263 – 29,670) than single-end libraries (18,526 – 40,651) (S1 Fig). In general, individual *de-novo* transcriptomes were of good quality and recovered 51-79% of the core eukaryotic universal single-copy orthologs [29] (S3 Table). The final *de novo* composite skin transcriptome reconstructed from all the RNA reads of the *Oophaga species* yielded 31,498 contigs greater than 350 bp. The overall assembly incorporated 94% of all initial reads and the level of fragmentation was low with half of all base pairs clustered into contigs of 1,316 bp in length or greater. The maximum contig length was 16,041 bp and the AL was 1014.91 ± 948.43 (S2A Fig). Nucleotide-based *BLAST* analyses (*BLASTX*) revealed that ~63% of the contigs (n=19,732) show significant similarities with either annotated gene products and/or known protein domains (E-value ≤ 10-5) (S2B Fig) and only a small fraction of unigenes (3.4%) showed significant homology to the same annotated transcript. The highest *BLAST* scores were found with *X. tropicalis* (74%), *X. laevis* (2%) and the green sea turtle *Chelonia mydas* (0.5%). The composite skin transcriptome was of higher quality when compared with individual assemblies, having 88.54% of the ortholog count of BUSCO [29] (S3 Table).

Gene Ontology (GO) assignments were used to classify the functions of the predicted unigenes based on contigs with significant *BLASTX* (E-value ≤ 10-5). Based on GO level II, unigenes were assigned to 27 biological processes (BP), 22 cell components (CC) and 15 molecular functions (MF) (S2C Fig). Some unigenes were associated with multiple GO annotations because a single sequence may be annotated in any or all categories, giving more GO annotations than sequences annotated [44]. Within BP, ~47% of the annotations were assigned to basic cellular, metabolic processes and biological regulation. The remaining unigenes were involved in a broad range of BP such as response to stimulus (7%), response to stress (7%), localization (6%), developmental process (5%), signal transduction (4%), biogenesis (4%), immune response (2%), reproductive process (1.7%) and cellular adhesion (1%). Within the CC category, other than the essential cell constituents (44%), the membrane components were highly represented in the transcriptome (19%). Within MF, most of the unigenes were assigned to binding and catalytic activities (72%) followed by transporter activity (9%), molecular transducers (7%) and molecular function regulators (4%) (S2C Fig).

### Transcriptome profiles

Dispersion plots of ER values in *Oophaga* RNA libraries showed a common pattern of over-represented unigenes across RNA-libraries. That is, the same group of unigenes showed similar patterns of over-representation no matter RNA library’s origin (*i*.*e*. different *Oophaga* species) or composition (paired *vs*. single-read libraries) (S3 Fig). The TPM represents a measure of RNA abundance and hence, it provides a general overview of gene expression levels in a particular sample. Interestingly, TPM distribution plots (percentiles) showed a distinct spike in expression levels towards percentile 98% for all our RNA-libraries (S4A Fig). After the conservative filtration of the 2% top expressed unigenes, we selected a subset of 1,437 unigenes with particular higher levels of expression (higher TPM values and PCA outliers, S4B Fig) in *Oophaga* skin tissue. Homology *BLAST* analysis of these unigenes revealed that 76% (n=1,092) show significant similarities with annotated gene products and/or known protein domains distributed mainly across amphibian (frogs) species (*X. tropicalis*=30%, *X. laevis*=14% and *Rana catesbeiana*=4%).

After adjusting for library size and length bias, the maximum effective number of reads (ER) in *Oophaga* species ranged from 2,461 (*O. anchicayensis* single-end) to 5,813 (*O. sylvatica* – single-end) while in the cryptic species varied from 82,509 (*Rana catesbeiana)* to 452,018 (*Bufo gargarizans*) (higher coverage in cryptic species, S2 Table). This indicated that after the optimization of our alignment parameters, a considerable amount of reads from each RNA-seq library actually mapped to our composite *de novo* transcriptome. After the removal of contigs with low expression profiles (<10 read counts), our RNA-seq dataset was composed of reads mapping to 20,771 genes. Multidimensional scaling plots (stress<0.05) revealed a clear separation by groups (*Oophaga* vs. cryptic species, p<0.001 Bonferroni corrected) with higher variation among the cryptic species, as expected given the broad taxonomic range of the members of this group (Fig 2).

**Fig 2.**
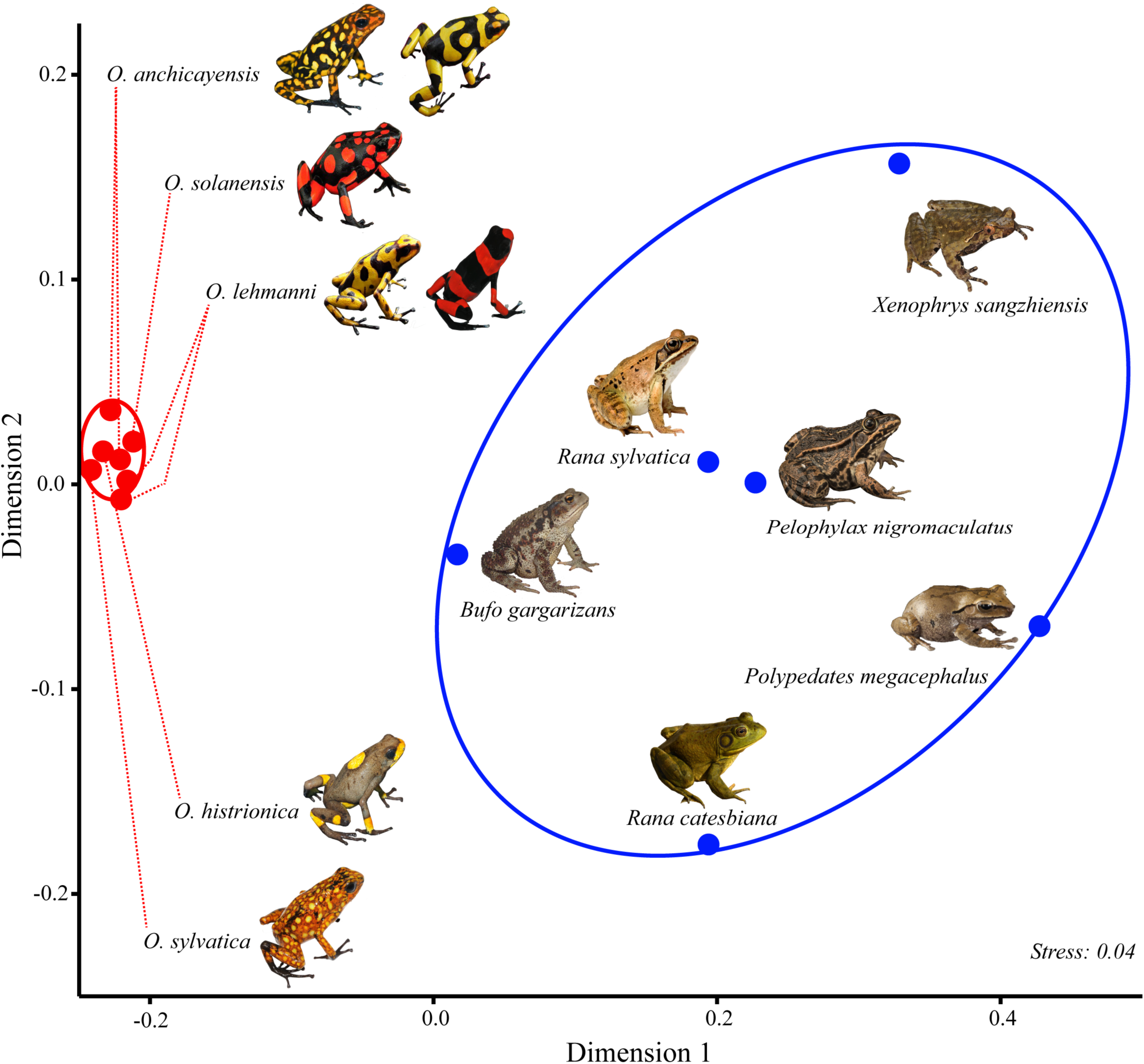
Multidimensional scaling (MDS) plot of 13 RNA-sequencing samples from skin tissue of *Oophaga* (n=7, red dots) and cryptic species (n=6, blue dots). RNA-seq data shown here represent the effective read counts (ER) for 31,498 skin-specific contigs. Color ellipses represent a 95% confidence interval.

About 9% of the highly expressed genes (n= 1,931) also showed a phylogenetic signal (*i*.*e*. significant fold changes in expression levels between *Oophaga* and all other anurans). Most of these genes (n=1,656; 85.8%) show higher levels of expression in *Oophaga*. The clustering analysis (Fig 1) revealed that the toad (*B. gargarizans)* has the closest transcription profile to *Oophaga* species. One possible explanation is the presence of alkaloid-based toxins in its skin glands [45].

Homology *BLAST* analyses revealed that 65% (n=1,252) show significant similarities with annotated gene products and/or known protein domains (S4 Table; S5A Fig) distributed mainly across amphibian (frogs) species (*X. tropicalis*=32%, *X. laevis*=19% and *Rana catesbeiana*=4%) (S5B Fig). Based on GO mapping level II, these unigenes were assigned to 24 BP, 20 CC and 13 MF (S5C Fig). Multilevel GO term classification assigned the highly expressed unigenes to 16 BP, 5 CC and 9 MF (S5C Fig). Within BP, 13% of unigenes were associated with oxidation-reduction processes, followed by response to stimulus (10%) and regulation of macromolecule metabolic processes (7.3%). Within CC, the integral components of the cell membrane were highly represented (39%), followed by protein complex (27%) and nuclear components (20%). Within MF, most of the differentially expressed unigenes were assigned to metal ion binding activity (21%), followed by protein binding activity and transmembrane transporters (17%), oxidoreductase activity (12%) and structural molecule activity (10%) (S5C Fig). The full functional annotation of these genes allowed us to propose plausible mechanisms that may be involved in the ability to sequester alkaloid-based chemical defenses, warning coloration and the auto-resistance mechanisms to avoid self-intoxication (S5 Table).

## Discussion

Studies focusing on highly represented and differential gene expression in related lineages have provided valuable insights into the molecular underpinnings of phenotypic variation [46, 47]. These analyses rely on the premises that lineages show contrasting phenotypes of interest, and that most differentially expressed genes relate to the trait of interest rather than the overall of genomic differentiation between lineages. Here, we examined shared patterns of gene expression in a small clade of closely related aposematic frogs (*Oophaga* species). Because all members of this monophyletic group show contrasting hues and accumulate alkaloid-derived toxins in their dermal glands [48, 49], we assumed that the genes related with these traits are likely to 1) be highly expressed in the skin of *Oophaga* poison frogs and 2) showed contrasting expression patterns between aposematic and cryptic and less (or not) poisonous species. Using expression levels in a multivariate framework, we first identified ortholog genes that are highly represented in *Oophaga* skin tissue and then, we compared with those differentially expressed in the skin of contrasting groups and that show similar patterns of expression among *Oophaga* species. We finally used a functional annotation analyses to highlight a series of genes that are possibly associated with a series of aposematic traits.

The characterization of our skin transcriptomic profiles allowed us to identify genes that are potentially associated with alkaloid transportation and resistance to autotoxicity in poison frogs (Dendrobatidae). Because the alkaloids secreted by these frogs are structurally very similar to those present in plants [18, 50–52], it is reasonable to assume that the transportation and accumulation of alkaloids in these frogs may be carried out by similar systems to those described in plants in which alkaloids are transferred from source to sink organs [53–55]. If so, at least, two alternative membrane mechanisms may help to explain the transport and storage of unmodified diet-derived alkaloid by the specialized cells of skin secretory glands. First, alkaloids in a lipophilic state may freely pass through the cell membranes by simple diffusion and accumulate in acidic secretory lysosomes if they become protonated to form hydrophilic cations. This ion-trap mechanism is not energy-dependent and does not necessarily require the expression of any transporters. Alternatively, the transportation of alkaloids may be managed by proton-antiport carrier systems in an energy-requiring manner [56–58]. Under this alternative model, diet derived alkaloids may be taken up by ABC transporters [59, 60] and may accumulate in the secretory lysosomes of the skin gland cells by a cation exchanger antiporter (CAX) system, dependent on the pH gradient generated by “vacuolar” type ATPases (V-H^+^-ATPases) and/or pyrophosphatases (V-H^+^-PPases). To find some support for these two potential models that might explain the accumulation and posterior secretion of dietary alkaloids, we investigated the common transcriptional profile across *Oophaga* species. For each of these species, our differential expression and transcript abundance analysis revealed that at least 15 homologs of different type II CAX are highly expressed in skin tissue (S5 Table). A result suggesting the existence of an active membrane transportation-accumulation mechanism of alkaloids in harlequin poison frogs. Consistent with this hypothesis, we also found putative ABC transporters and at least one V-H^+^-ATPase on the skin transcriptome. All together, these results suggest that active lysosomal exocytosis may play a key role in the secretion of alkaloids in these frogs.

The accumulation of toxic compounds implies that organisms must avoid self-intoxication (auto-resistance). While the membrane transportation mechanisms described above reduce auto-toxicity by compartmentalizing the sequestered alkaloids, other non-alternative excluding mechanisms are likely to contribute to auto-toxicity resistance. In harlequin poison frogs, the major toxic components present in skin tissue are histrionicoxins (HTX) [51]. These alkaloids are known to cause temporary paralysis or even death by inactivating or blocking voltage-gate ion channels [61]. Resistance to this type of cytotoxic compounds usually arises through either, an increased expression level of P450 enzymes (CYPs) that metabolize the toxin, or through target insensitivity via mutations that reduce the toxin’s ability to bind the ion channel itself [62]. Our analyses revealed that five CYP homologs are amongst the most highly expressed genes in the skin of harlequin poison frogs (S5 Table). This result suggests that the oxidative biotransformation of lipophilic alkaloids to hydrophilic compounds [63] is an important auto resistance mechanism in these frogs. Although the overproduction of the CYP-enzymes has been interpreted as evidence for metabolic detoxification [64], there is an alternative explanation for the high expression levels of CYP proteins in skin of harlequin poison frogs. While many of the alkaloids sequestered in the wild are accumulated unchanged in the dermal glands, at least two species of dendrobatids, stereo-selectively hydroxylate the toxic pumiliotoxin PTX-(+)-251D to convert it into a much more potent toxin, the allopumiliotoxin aPTX-(+)-267A [17]. Both PTX-(+)-251D and aPTX-(+)-267A have been isolated from the skin of *O. histrionica*, and CYPs are known to be involved in the stereo-selective hydroxylation of alkaloids [65]. Thus, it is also possible that the CYPs expressed in the skin of harlequin poison frogs may enhance the antipredator potency of ingested PTXs.

Despite the major role that detoxification enzymes may have in the resistance to diet acquired toxins, mutations conferring constitutive resistance to alkaloids are also likely to be involved in auto-resistance in chemically-defended dendrobatids. In insects, single amino-acid substitution in the voltage-gated sodium channels (Na^+^K^+^-ATPases) are responsible for resistance to host-plant phytochemicals [66–68]. In amphibians, the only documented example of this this type of target insensitivity is from distantly-related lineages of alkaloid-defended frogs, 24 species of Neotropical dendrobatids and the aposematic Madagascar frog *Mantella aurantiaca* [69]. In all of these cases, they carried different amino acid replacements in the inner pore of the voltage-gated sodium channel Nav1.4 that are not found in other frog [70]. Our analyses revealed eight genes encoding voltage-gated ion channel proteins (VGIC), one of which (8966), encodes the gamma-1 subunit of a sodium channel.

Aposematism in harlequin frogs is a complex phenotype that results from the integration of different elements including unpalatability and conspicuousness. Thus, another of our goals was to identify genes potentially related with warning coloration. Many aposematic organisms, including the *Oophaga* poison frogs studied here exhibit color patterns that show strong color and/or luminance contrast such between black and red, orange or yellow [15]. It has been proposed that such patterning increases aversion learning as well as conspicuousness and distinctiveness from palatable prey [71–76]. In anurans, coloration relates to both the structure and pigment composition of the dermal chromatophore units in which three types of pigment cells (xanthophores, iridophores and melanophores) are laid one on another [77]. In dark/black skin areas of harlequin poison frogs, no xanthophores or iridophores are present and the distal fingers of the melanophores are filled with melanin granules (melanosomes) obscuring the dermis [15]. Thus, genes involved in the amount, size and distribution of the melanosomes are likely to play a significant role in the coloration pattern of these frogs.

In one of the few detailed studies of coloration in anurans [78], authors suggest a common origin for of all the pigment granules found in the cells of the chromatophore: a primordial organelle (vesicle) derived from the rough endoplasmic reticulum (RER). According to this model, in the formation of melanosomes, the pre-melanosomes are derived from cisternae of the RER, which then fuse with vesicles derived from the Golgi complex containing tyrosinase enzymes [12, 78–81]. A highly expressed gene identified here (tyrosinase regulator, 25062) may contribute to regulate this mechanism, which in turn might translate into hue differences. Interestingly, among *Oophaga* species, we have found populations that are characterized by light-brown background coloration as oppose to black [15, 22]. One tantalizing possibility is that this phenotypic difference is indeed associated with the expression differences of these tyrosinase enzymes.

Obviously, there are other molecular and cellular mechanisms that might be associated to the difference between light and dark background coloration. In fishes and amphibians, dark hues are known to be produced by the interaction between high levels of melanocyte-stimulating hormone (α-MSH) and several variants of its transmembrane receptor (*MC1R*) through the dispersion of melanosomes within the melanophore (by increasing cAMP intracellular levels) [82, 83]. Thus, it is possible that structural or expression differences in the *MC1R* might contribute to dark phenotypes in *Oophaga* frogs. While there is a strong evidence that different mutations at *MC1R* cause either light or dark phenotypes in many mammals, birds and reptiles [84–88], the only two studies conducted in frogs are inconclusive [89, 90]. A detailed inspection of the coding sequences recovered for this gene revealed the presence of different length isoforms, making *MC1R* a promising candidate gene candidate to explain the differences in background coloration in poison frogs [15]. Finally, our study revealed another two highly and differentially expressed genes (14447-Melanoregulin-; 2038-Melanocortin phosphoprotein), which may also contribute to dark hues. In this case, the predicted products of these genes are key proteins that mediate the melanosome transport and distribution in epidermal cells through the formation of a tripartite protein complex [91]. The disruption of the transport protein complex results in pigmentary dilution and lighter phenotypes by the clustering of melanosomes around the nucleus [92].

Overall, this study demonstrates the utility of using RNA-sequencing with non-model organisms to identify loci that may be of adaptive importance. Altogether, these data enabled us to provide a first global study of *Oophaga* poison frogs transcriptomes and propose potential mechanisms for alkaloid sequestration, auto-resistance to toxic compounds, and variation in coloration and patterns. It is important to recognized that due the sampling restrictions imposed to us (see methods), we are most likely missing some of the genes responsible for these traits. It is also probable that the genes associated with coloration, alkaloid metabolism, transport and storage are differentially expressed not only in skin, but other tissues in the organism. Despite the potential caveats, other molecular studies have shown that some of the genes identified in this study are indeed potentially related with aposematic traits in dendrobatids [15, 70, 93].

Despite the critical impact of genetic basis of coloration in the evolution and diversification of aposematic phenotypes, the genetic architecture of coloration in poison frogs (Dendrobatidae) remains virtually unexplored. Here, we report probably the first RNA-seq study in *Oophaga* poison frogs, a model system for understanding the relationship between toxicity, diet and coloration. The skin-expressed genes that we have identified here provide initial working hypotheses to further unravel the molecular genetics mechanisms to sequester alkaloid-based chemical defenses, warning coloration and the auto-resistance mechanisms to avoid self-intoxication. Hence, further analysis aiming to compare the amino acid substitutions in the expressed cDNAs will provide insights about the structure and function of each of these genes. Finally, our comparative transcriptome data provide an important new resource to better understand the evolution of warning signals in nature.

## Acknowledgements

We would like to thank the former members of Rick Harrison’s lab (*requiescat in pace*) at Cornell University for their valuable discussions and comments on early versions of the manuscript, which was greatly improved by the thoughtful comments of Dr. Andrew Crawford at Universidad de los Andes (Colombia). This study was funded by COLCIENCIAS (Departamento Administrativo de Ciencia, Tecnología e Innovación, Colombia, 529-2011) grant to A.P.-T. and an NSERC-Discovery grant to J.A.A.

## Supporting information

### Supporting Tables

**S1 Table.** Number of reads and library type (i.e paired vs. single-end) for each individual RNA-seq experiment.

| Individual library | Raw number of reads | Reads after filtering | Remaining % of reads | Library type |
| --- | --- | --- | --- | --- |
| *O. solanensis* | 128,599,724 | 105,179,065 | 81% | Paired reads* |
| *O. anchicayensis* | 124,232,804 | 110,217,128 | 88% |  |
| *O. lehmanni* | 102,225,680 | 91,493,848 | 89% |  |
| *O. sylvatica* | 48,929,329 | 45,147,654 | 92% | Single reads |
| *O. lehmanni* | 43,633,080 | 41,086,379 | 94% |  |
| *O. anchicayensis*) | 7,998,636 | 7,173,285 | 89% |  |
| *O. histrionica* | 50,204,617 | 47,376,912 | 94% |  |
| Reference transcriptome | N/A | 33,895,474 | 100% | Single reads |

\*. Number of reads correspond to the combined value of both ends libraries.

**S2 Table.** RNA-sequence datasets from cryptic species used for the differential expression analysis of this work.

| Species | Experiment (SRA*) | RNA-seq library | Total bases | Date available |
| --- | --- | --- | --- | --- |
| *Bufo gargarizans* | SRX2640691 | Paired-end (150 b) | 19.2 Gbp | 2017-03-15 |
| *Pelophylax nigromaculatus* | SRX2640690 | Paired-end (150 b) | 18.0 Gbp | 2017-03-15 |
| *Polypedates megacephalus* | SRX1720193 | Paired-end (100 b) | 6.3 Gbp | 2016-04-25 |
| *Rana catesbeiana* | SRX2988982 | Paired-end (100 b) | 9.2 Gbp | 2017-07-07 |
| *Rana sylvatica* | SRX2989023 | Paired-end (100 b) | 8.9 Gbp | 2017-07-07 |
| *Xenophrys sangzhiensis* | SRX1720191 | Paired-end (100 b) | 6.8 Gbp | 2016-04-25 |

\* Sequence Read Archive accession number from Genbank.

**S3 Table.** Quality assessment report for each individual and the composite reference transcriptome based on the protein count of the core eukaryotic genes (BUSCO, see methods in the main article)

| Transcriptome | Library | Busco v.2 mapping |  |  |
| --- | --- | --- | --- | --- |
|  |  | Complete | Partial | Complete + Partial |
| *O. solanensis* | Paired | 20.60% | 30.47% | 51.07% |
| *O. anchicayensis* | Paired | 24.46% | 37.77% | 62.23% |
| *O. lehmanni* | Paired | 29.61% | 40.77% | 70.38% |
| *O. sylvatica* | Single | 45.92% | 33.48% | 79.40% |
| *O. lehmanni* | Single | 33.91% | 39.91% | 73.82% |
| *O. anchicayensis* | Single | 9.87% | 30.47% | 40.34% |
| *O. histrionica* | Single | 39.06% | 37.77% | 76.83% |
| Reference assembly | Single | 45.92% | 42.62% | 88.54% |

**S4 Table.** Summary statistics of the differentially expressed unigenes (see methods) and their *BLAST* annotations.

|  | Differentially expressed unigenes | With annotation | Without annotation |
| --- | --- | --- | --- |
| **Up-regulated** | 1656 (85.6%) | 996 (51.6%) | 660 (34.2%) |
| **Down-regulated** | 275 (14.2%) | 256 (13.3%) | 19 (1%) |
| **Total** | 1931 (100%) | 1252 (64.9%) | 679 (35.2%) |

**S5 Table.**
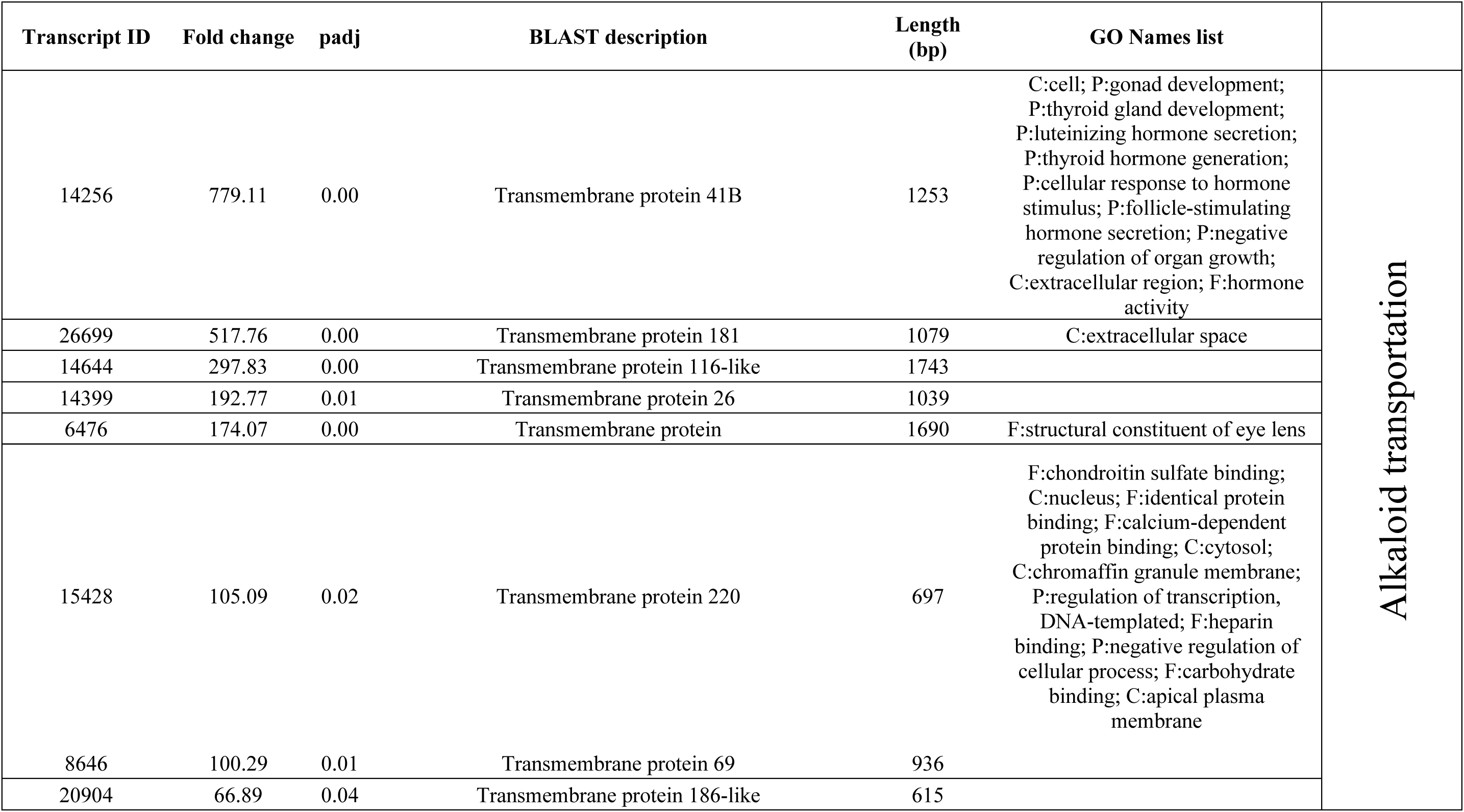
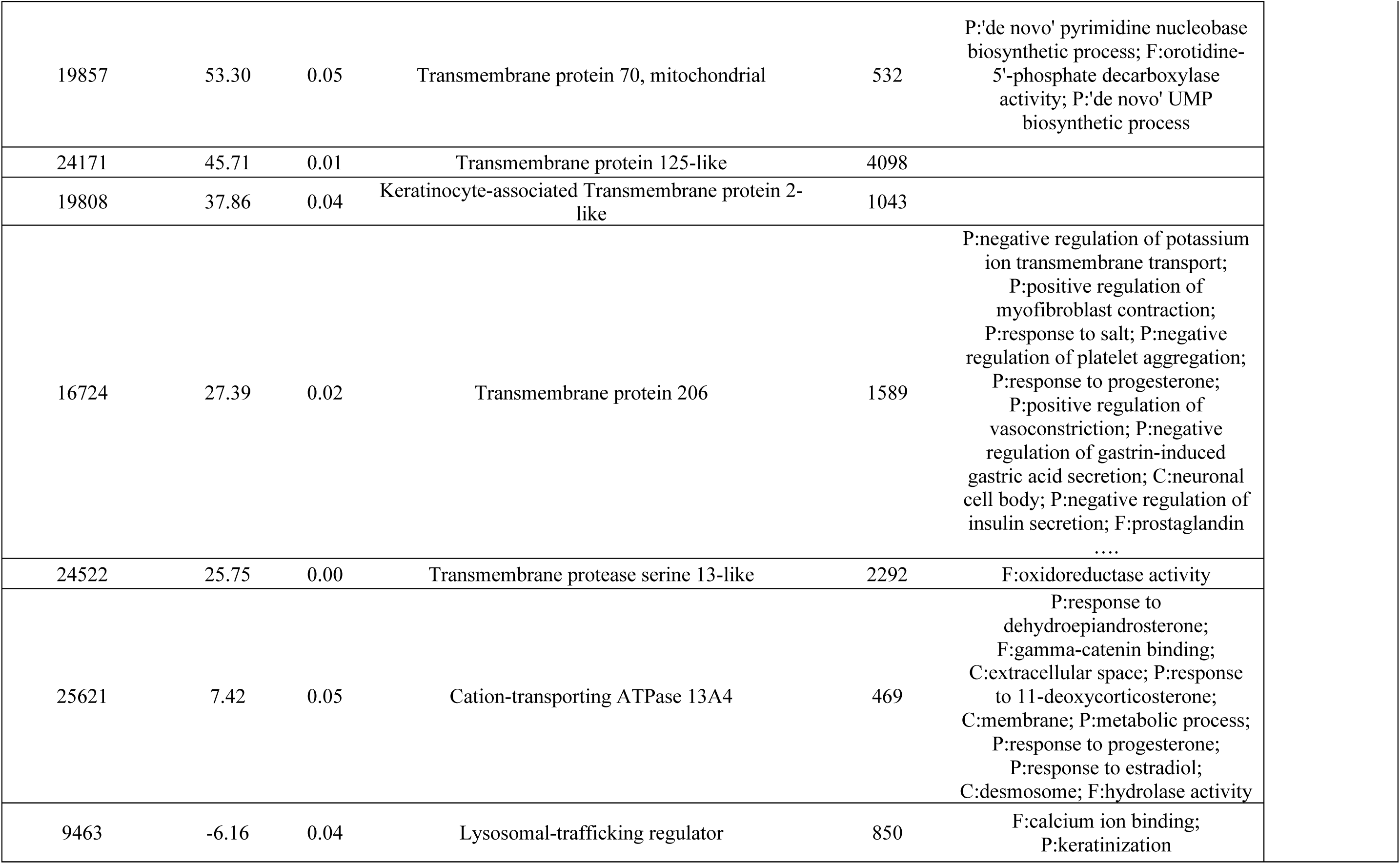
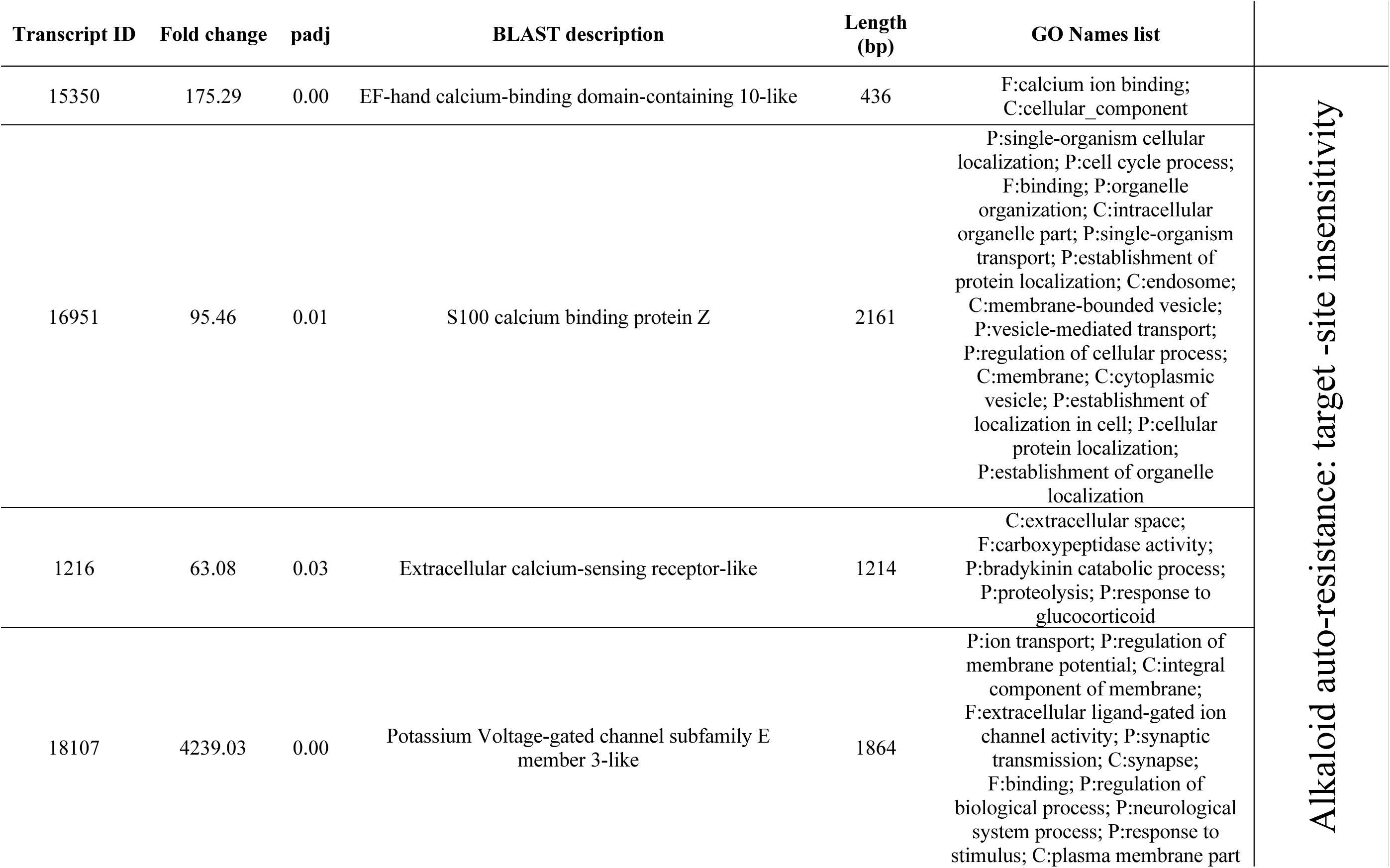
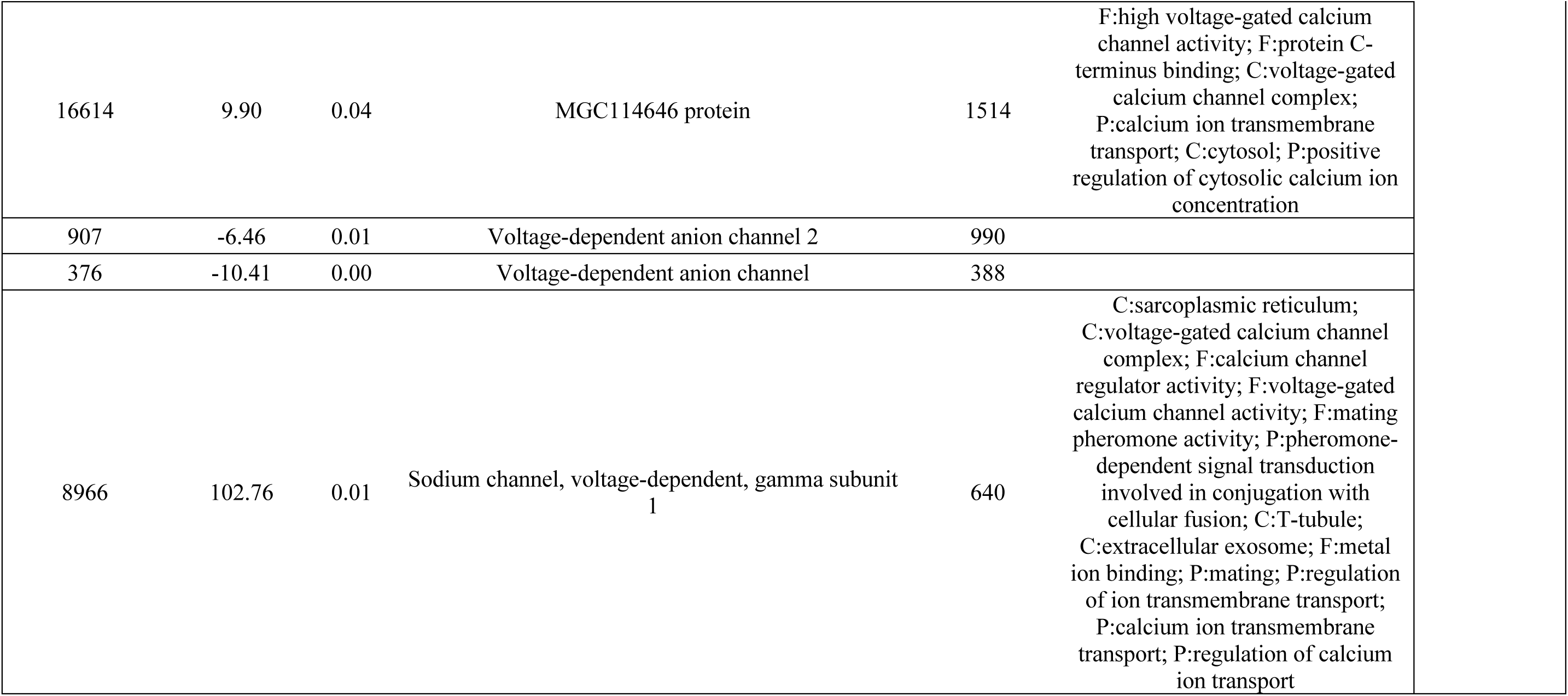
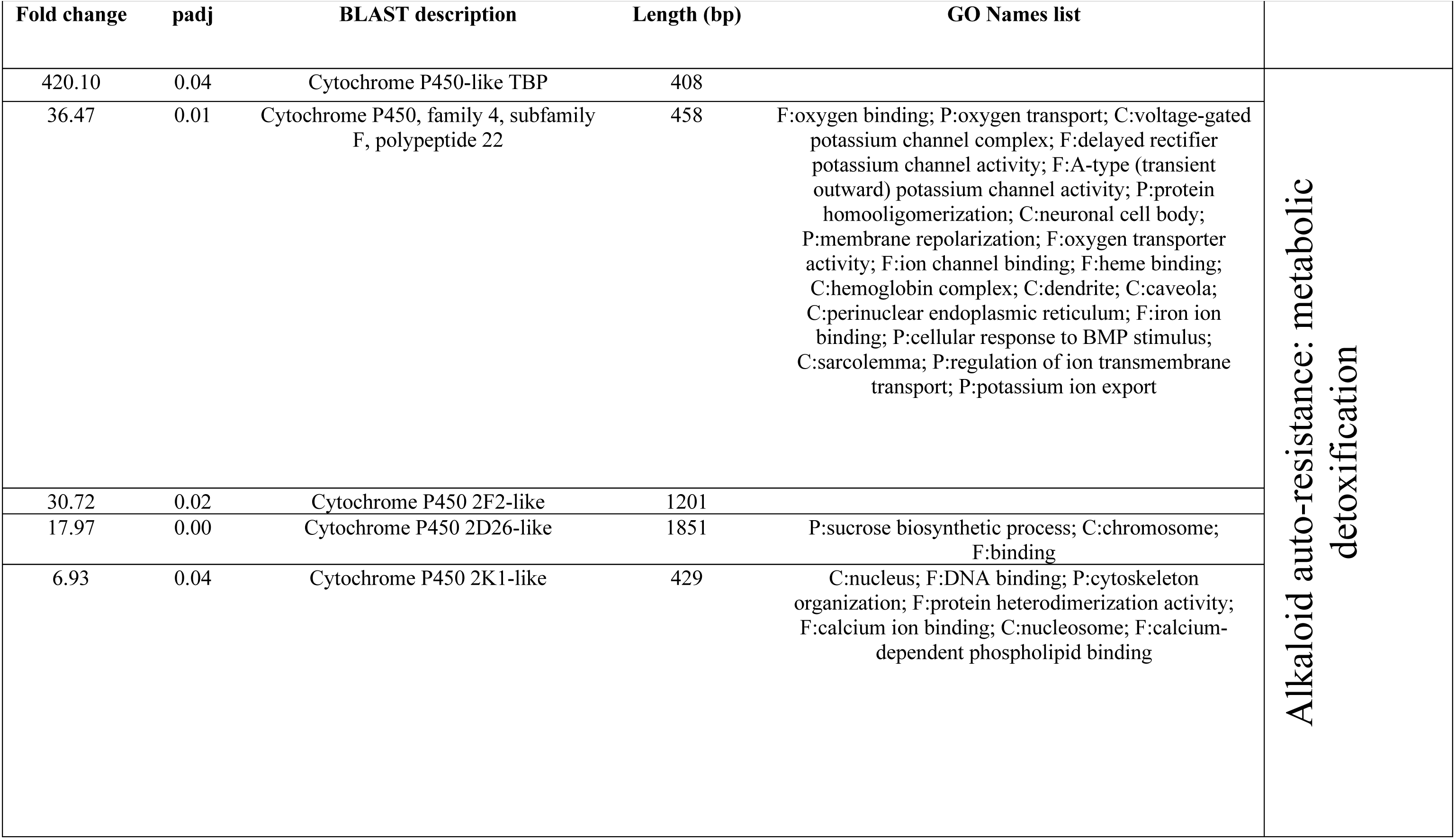
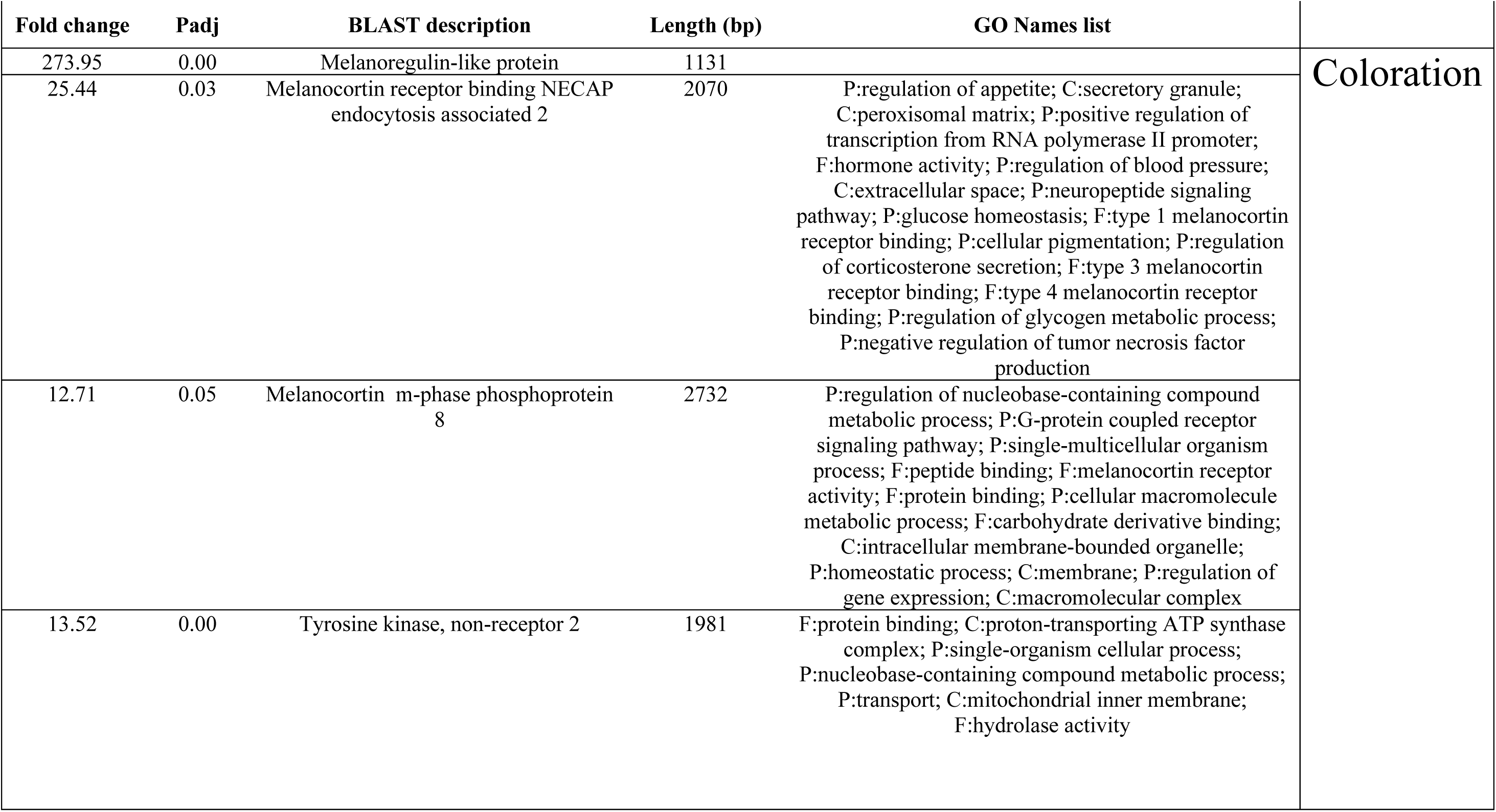
Transcripts with potential important functions in the alkaloid sequestration system, auto-resistance to toxic compounds and variation in coloration in harlequin poison frogs. The corresponding composite *Oophaga* transcriptome is available from the authors upon request.

| Transcript ID | Fold change | padj | BLAST description | Length (bp) | GO Names list |  |
| --- | --- | --- | --- | --- | --- | --- |
| 14256 | 779.11 | 0.00 | Transmembrane protein 41B | 1253 | C:cell; P:gonad development; P:thyroid gland development; P:luteinizing hormone secretion; P:thyroid hormone generation; P:cellular response to hormone stimulus; P:follicle-stimulating hormone secretion; P:negative regulation of organ growth; C:extracellular region; F:hormone activity | Alkaloid transportation |
| 26699 | 517.76 | 0.00 | Transmembrane protein 181 | 1079 | C:extracellular space |  |
| 14644 | 297.83 | 0.00 | Transmembrane protein 116-like | 1743 |  |  |
| 14399 | 192.77 | 0.01 | Transmembrane protein 26 | 1039 |  |  |
| 6476 | 174.07 | 0.00 | Transmembrane protein | 1690 | F:structural constituent of eye lens |  |
| 15428 | 105.09 | 0.02 | Transmembrane protein 220 | 697 | F:chondroitin sulfate binding; C:nucleus; F:identical protein binding; F:calcium-dependent protein binding; C:cytosol; C:chromaffin granule membrane; P:regulation of transcription, DNA-templated; F:heparin binding; P:negative regulation of cellular process; F:carbohydrate binding; C:apical plasma membrane |  |
| 8646 | 100.29 | 0.01 | Transmembrane protein 69 | 936 |  |  |
| 20904 | 66.89 | 0.04 | Transmembrane protein 186-like | 615 |  |  |

|  |  |  |  |  |  |
| --- | --- | --- | --- | --- | --- |
| 19857 | 53.30 | 0.05 | Transmembrane protein 70, mitochondrial | 532 | P:'de novo' pyrimidine nucleobase biosynthetic process; F:orotidine-5'-phosphate decarboxylase activity; P:'de novo' UMP biosynthetic process |
| 24171 | 45.71 | 0.01 | Transmembrane protein 125-like | 4098 |  |
| 19808 | 37.86 | 0.04 | Keratinocyte-associated Transmembrane protein 2-like | 1043 |  |
| 16724 | 27.39 | 0.02 | Transmembrane protein 206 | 1589 | P:negative regulation of potassium ion transmembrane transport; P:positive regulation of myofibroblast contraction; P:response to salt; P:negative regulation of platelet aggregation; P:response to progesterone; P:positive regulation of vasoconstriction; P:negative regulation of gastrin-induced gastric acid secretion; C:neuronal cell body; P:negative regulation of insulin secretion; F:prostaglandin .... |
| 24522 | 25.75 | 0.00 | Transmembrane protease serine 13-like | 2292 | F:oxidoreductase activity |
| 25621 | 7.42 | 0.05 | Cation-transporting ATPase 13A4 | 469 | P:response to dehydroepiandrosterone; F:gamma-catenin binding; C:extracellular space; P:response to 11-deoxycorticosterone; C:membrane; P:metabolic process; P:response to progesterone; P:response to estradiol; C:desmosome; F:hydrolase activity |
| 9463 | -6.16 | 0.04 | Lysosomal-trafficking regulator | 850 | F:calcium ion binding; P:keratinization |
| 15350 | 175.29 | 0.00 | EF-hand calcium-binding domain-containing 10-like | 436 | F:calcium ion binding;<br>C:cellular_component | Alkaloid auto-resistance: target -site insensitivity |
| 16951 | 95.46 | 0.01 | S100 calcium binding protein Z | 2161 | P:single-organism cellular localization; P:cell cycle process; F:binding; P:organelle organization; C:intracellular organelle part; P:single-organism transport; P:establishment of protein localization; C:endosome; C:membrane-bounded vesicle; P:vesicle-mediated transport; P:regulation of cellular process; C:membrane; C:cytoplasmic vesicle; P:establishment of localization in cell; P:cellular protein localization; P:establishment of organelle localization |  |
| 1216 | 63.08 | 0.03 | Extracellular calcium-sensing receptor-like | 1214 | C:extracellular space; F:carboxypeptidase activity; P:bradykinin catabolic process; P:proteolysis; P:response to glucocorticoid |  |
| 18107 | 4239.03 | 0.00 | Potassium Voltage-gated channel subfamily E member 3-like | 1864 | P:ion transport; P:regulation of membrane potential; C:integral component of membrane; F:extracellular ligand-gated ion channel activity; P:synaptic transmission; C:synapse; F:binding; P:regulation of biological process; P:neurological system process; P:response to stimulus; C:plasma membrane part |  |

|  |  |  |  |  |  |
| --- | --- | --- | --- | --- | --- |
| 16614 | 9.90 | 0.04 | MGC114646 protein | 1514 | F:high voltage-gated calcium channel activity; F:protein C-terminus binding; C:voltage-gated calcium channel complex; P:calcium ion transmembrane transport; C:cytosol; P:positive regulation of cytosolic calcium ion concentration |
| 907 | -6.46 | 0.01 | Voltage-dependent anion channel 2 | 990 |  |
| 376 | -10.41 | 0.00 | Voltage-dependent anion channel | 388 |  |
| 8966 | 102.76 | 0.01 | Sodium channel, voltage-dependent, gamma subunit 1 | 640 | C:sarcoplasmic reticulum; C:voltage-gated calcium channel complex; F:calcium channel regulator activity; F:voltage-gated calcium channel activity; F:mating pheromone activity; P:pheromone-dependent signal transduction involved in conjugation with cellular fusion; C:T-tubule; C:extracellular exosome; F:metal ion binding; P:mating; P:regulation of ion transmembrane transport; P:calcium ion transmembrane transport; P:regulation of calcium ion transport |

| Fold change | padj | BLAST description | Length (bp) | GO Names list |  |
| --- | --- | --- | --- | --- | --- |
| 420.10 | 0.04 | Cytochrome P450-like TBP | 408 |  | Alkaloid auto-resistance: metabolic detoxification |
| 36.47 | 0.01 | Cytochrome P450, family 4, subfamily F, polypeptide 22 | 458 | F:oxygen binding; P:oxygen transport; C:voltage-gated potassium channel complex; F:delayed rectifier potassium channel activity; F:A-type (transient outward) potassium channel activity; P:protein homooligomerization; C:neuronal cell body; P:membrane repolarization; F:oxygen transporter activity; F:ion channel binding; F:heme binding; C:hemoglobin complex; C:dendrite; C:caveola; C:perinuclear endoplasmic reticulum; F:iron ion binding; P:cellular response to BMP stimulus; C:sarcolemma; P:regulation of ion transmembrane transport; P:potassium ion export |  |
| 30.72 | 0.02 | Cytochrome P450 2F2-like | 1201 |  |  |
| 17.97 | 0.00 | Cytochrome P450 2D26-like | 1851 | P:sucrose biosynthetic process; C:chromosome; F:binding |  |
| 6.93 | 0.04 | Cytochrome P450 2K1-like | 429 | C:nucleus; F:DNA binding; P:cytoskeleton organization; F:protein heterodimerization activity; F:calcium ion binding; C:nucleosome; F:calcium-dependent phospholipid binding |  |

| Fold change | Padj | BLAST description | Length (bp) | GO Names list | Coloration |
| --- | --- | --- | --- | --- | --- |
| 273.95 | 0.00 | Melanoregulin-like protein | 1131 |  |  |
| 25.44 | 0.03 | Melanocortin receptor binding NECAP endocytosis associated 2 | 2070 | P:regulation of appetite; C:secretory granule; C:peroxisomal matrix; P:positive regulation of transcription from RNA polymerase II promoter; F:hormone activity; P:regulation of blood pressure; C:extracellular space; P:neuropeptide signaling pathway; P:glucose homeostasis; F:type 1 melanocortin receptor binding; P:cellular pigmentation; P:regulation of corticosterone secretion; F:type 3 melanocortin receptor binding; F:type 4 melanocortin receptor binding; P:regulation of glycogen metabolic process; P:negative regulation of tumor necrosis factor production |  |
| 12.71 | 0.05 | Melanocortin m-phase phosphoprotein 8 | 2732 | P:regulation of nucleobase-containing compound metabolic process; P:G-protein coupled receptor signaling pathway; P:single-multicellular organism process; F:peptide binding; F:melanocortin receptor activity; F:protein binding; P:cellular macromolecule metabolic process; F:carbohydrate derivative binding; C:intracellular membrane-bounded organelle; P:homeostatic process; C:membrane; P:regulation of gene expression; C:macromolecular complex |  |
| 13.52 | 0.00 | Tyrosine kinase, non-receptor 2 | 1981 | F:protein binding; C:proton-transporting ATP synthase complex; P:single-organism cellular process; P:nucleobase-containing compound metabolic process; P:transport; C:mitochondrial inner membrane; F:hydrolase activity |  |

### Supporting figures

**S1 Fig.**
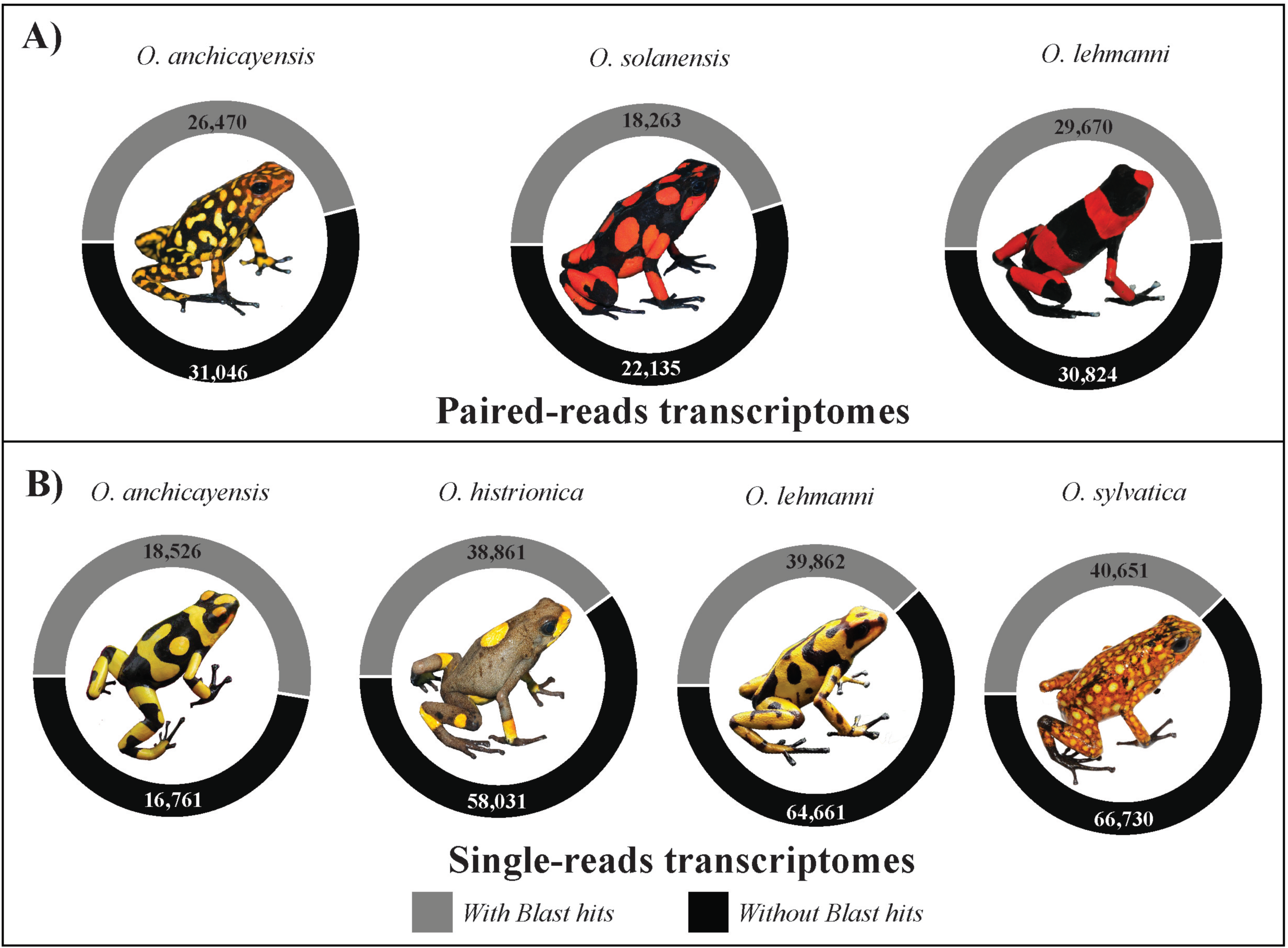
Pie charts representing the number and proportion of contigs with significant *BLAST* hits (E<1.0E^−5^) for each individual transcriptome from *Oophaga* species. Transcriptomes obtained using paired-ends RNA-seq libraries are represented in (A) and those obtained with single-end libraries in (A).

**S2 Fig.**
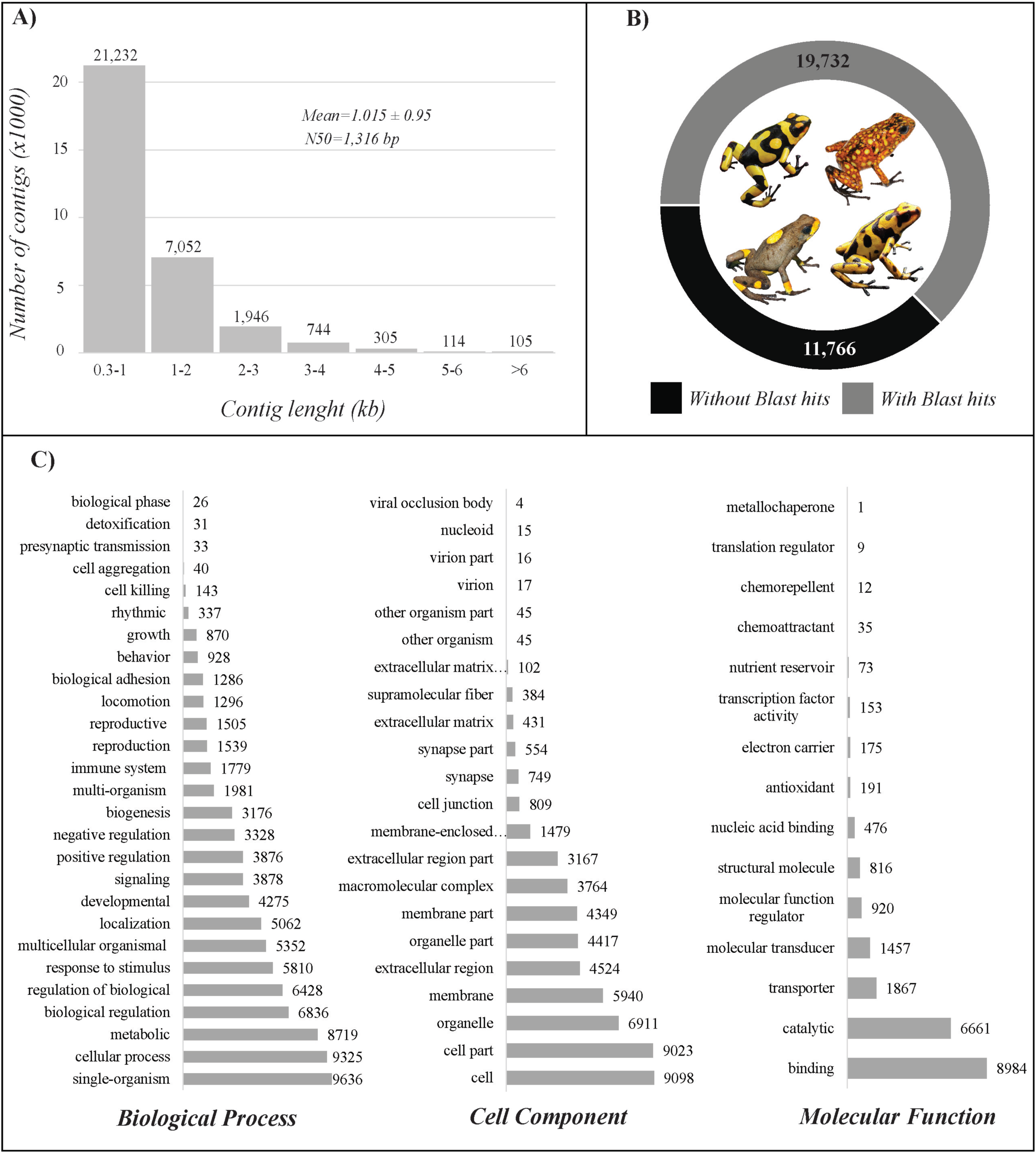
A) Contig length distribution of the *de-novo* composite reference transcriptome of *Oophaga* species from skin tissue. B) Pie chart representing the number and proportion of contigs with significant *BLAST* hits (E<1.0E^−5^) in the reference *Oophaga* transcriptome. C) Gene ontology (GO) categories distribution (level II) for the annotated unigenes in the *Oophaga* reference transcriptome.

**S3 Fig.**
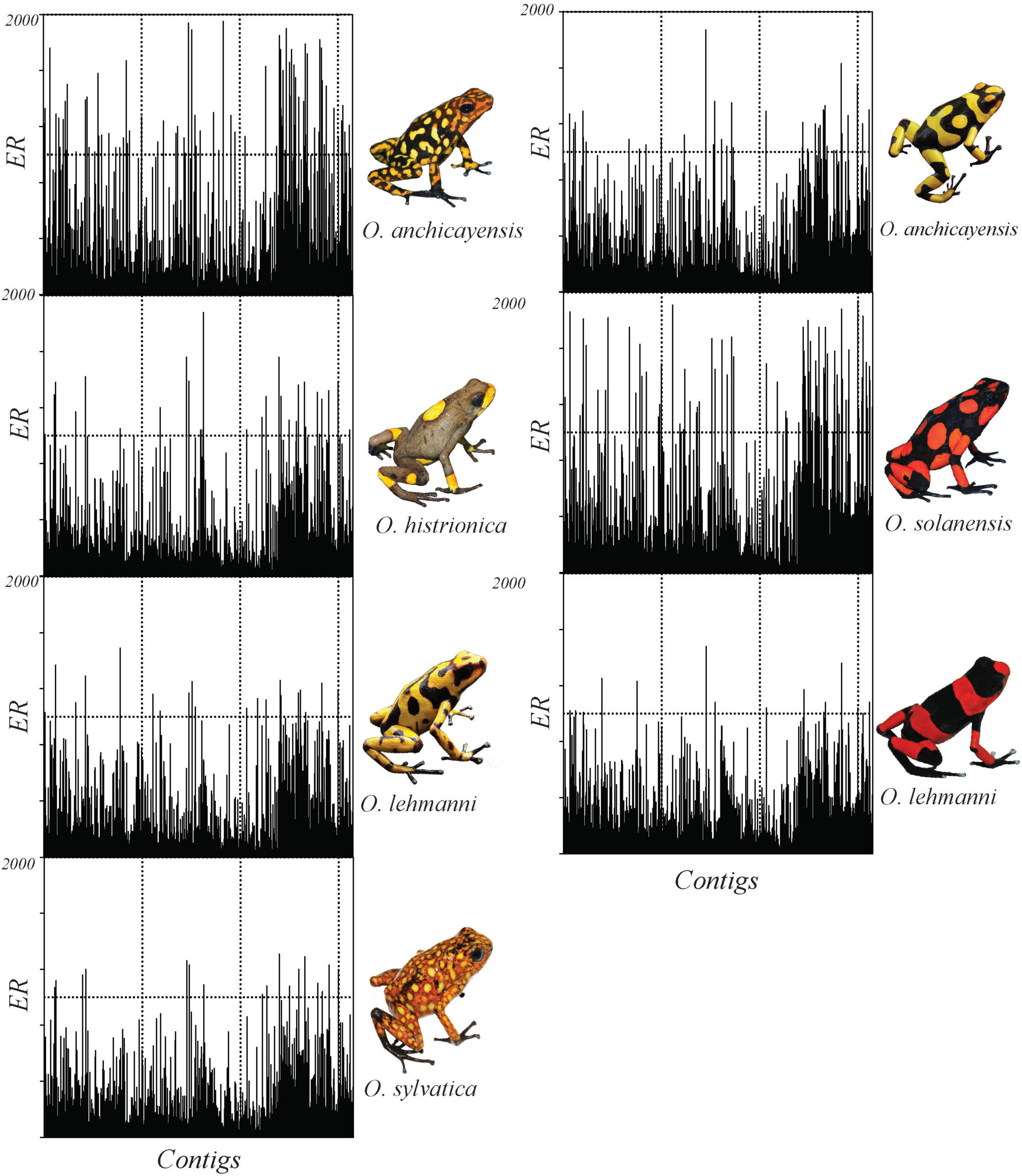
Distribution plots of the effective number of reads (ER) from *Oophaga* RNA-seq experiments that mapped to the composite reference transcriptome contigs (n=31,498).

**S4 Fig.**
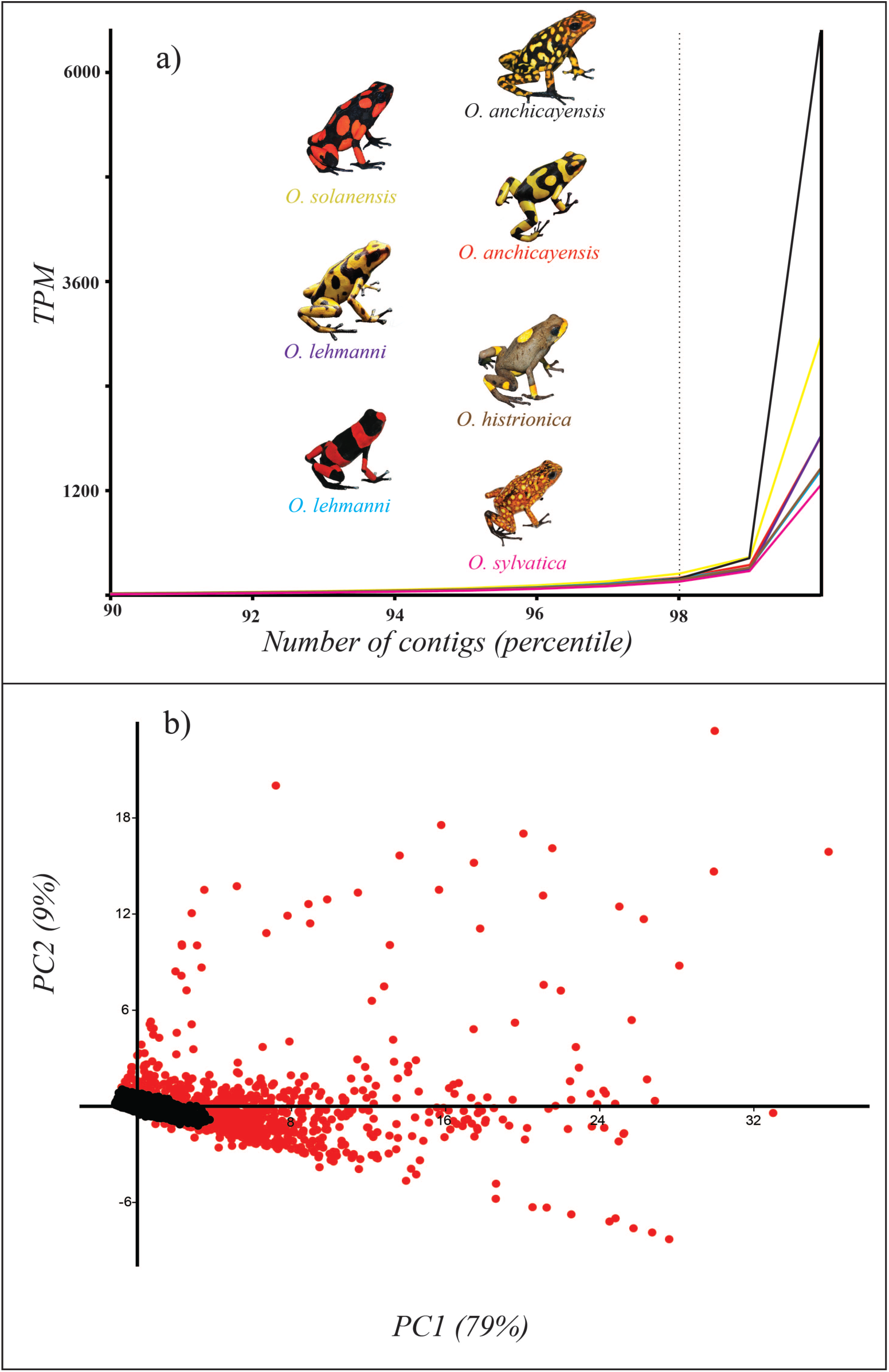
a) Percentile plot of the estimated transcripts per million (TPM) in the composite reference transcriptome as calculated based on raw RNA reads from individual libraries. Color names are equivalent to those in tendency lines. b) Principal component analyses (PCA) plot of the reference contigs dataset using TPMs values as independent variables. Red dots represent the selected highly represented unigenes (2% of the total contigs; n= 1,437) while dark dots represent the remaining ones (n=30,061)

**S5 Fig.**
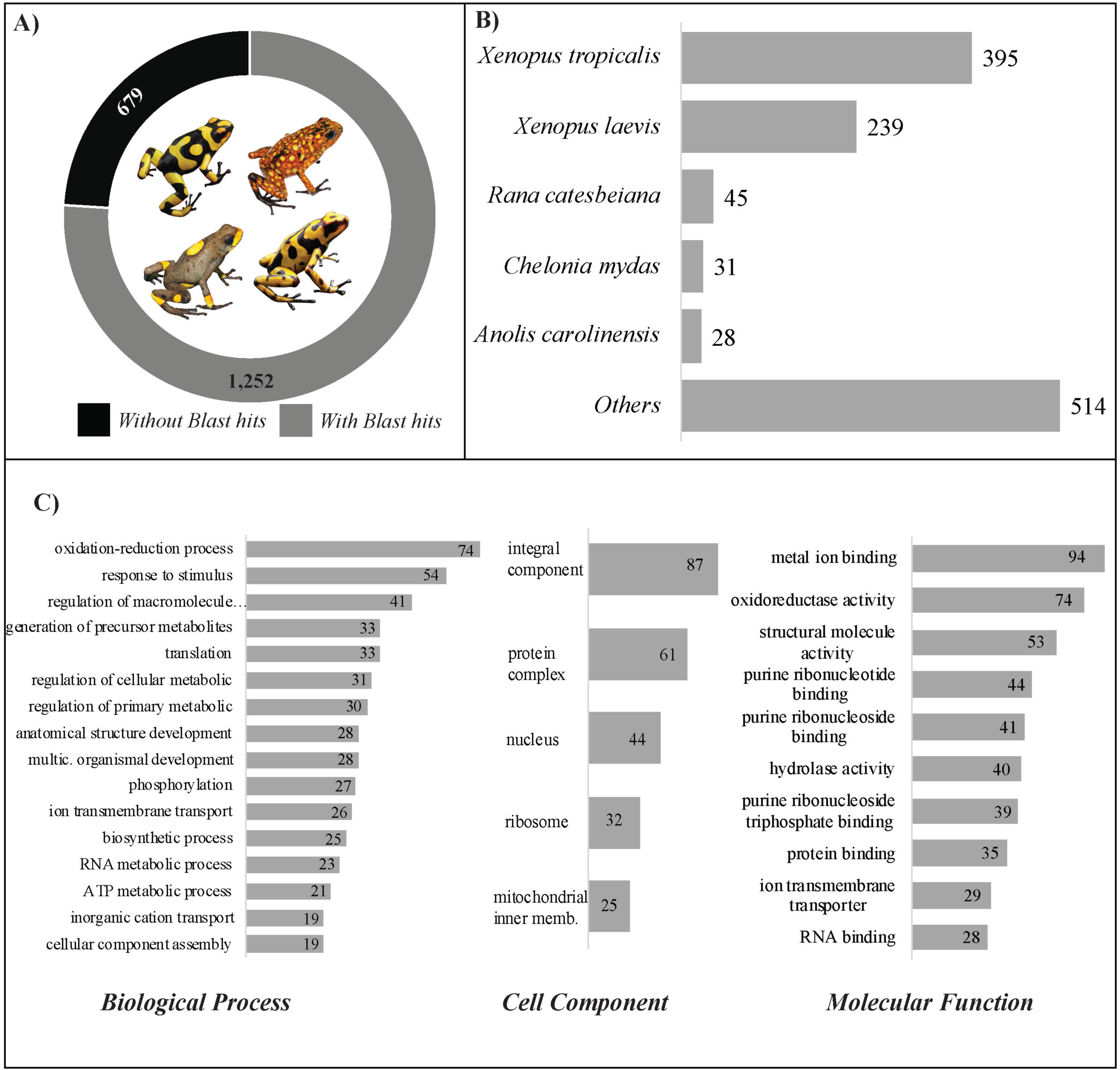
A) Pie chart representing the total number of differentially expressed unigenes and the proportion with significant *BLAST* hits (E<1.0E^−5^). B) BLAST hit species distribution of highly expressed unigenes. C) Gene ontology (GO) categories distribution (multilevel) for the annotated highly expressed unigenes.

